# Parallel occurrence of theta and respiration-coupled network oscillations throughout the mouse brain

**DOI:** 10.1101/139485

**Authors:** Adriano BL Tort, Simon Ponsel, Jakob Jessberger, Yevgenij Yanovsky, Jurij Brankačk, Andreas Draguhn

## Abstract

Slow brain oscillations are usually coherent over long distances and are thought to constitute a means to link distributed cell assemblies. In mice, theta oscillations (4-12 Hz) stand as one of the most studied global slow rhythms. Previous research has suggested that theta takes part in interregional communication required for cognitive functions. However, mice often breathe at theta frequency, and we have recently reported that nasal respiration leads to synchronous network oscillations that are independent of theta. Namely, we showed that respiration-coupled oscillations occur in the hippocampus, prelimbic cortex, and parietal cortex, suggesting that, as theta, respiration-coupled oscillations are also global. In the present work, we sought to extend these findings by tracking respiration while simultaneously recording local field potentials from 15 brain regions of freely moving mice during exploration and REM sleep. We report that respiration-coupled rhythms can be detected in parallel with theta in widespread neocortical regions, from prefrontal to visual areas, and also in subcortical structures such as the thalamus, amygdala and ventral hippocampus. Though both rhythms occur simultaneously, respiration-coupled oscillations are more dominant in frontal regions while theta oscillations prevail in more caudal networks. We conclude that respiration-coupled oscillations are a global brain rhythm suited to entrain distributed networks into a common regime. This pattern might have escaped attention in previous studies due to the absence of respiration monitoring, its similarity with theta oscillations, and its highly variable peak frequency. It should, however, be considered as a widespread signal and potential mechanism of long-range network communication.

## Introduction

Oscillations are ubiquitous in the electrical activity produced by the brain (Buzsáki & Draguhn, 2004). They can be observed at multiple scales, from spike times of single neurons, through the mesoscopic scale of local field potentials (LFPs), up to more macroscopic EEG and fMRI recordings (Buzsáki *et al.*, 2012). Though brain oscillations are usually classified according to their frequency, other factors influence their definition, such as wave shape, recorded region, animal species, and associated behavior (Wang, 2010; Cole & Voytek, 2017). These parameters are important because different types of oscillatory activity can occur within the same frequency range (Tort *et al.*, 2010; Lockmann *et al.*, 2016). Ideally, oscillations should be classified based on their origin and physiological function. Unfortunately, such a classification is currently not possible since, for most oscillations, we still do not have a full understanding of their underlying mechanisms.

We have recently studied a new type of LFP oscillations in rats and mice that fail to conform to a strict frequency-based definition: the so-called respiration-entrained rhythm (RR) (Yanovsky *et al.*, 2014; Jessberger *et al.*, 2016; Lockmann *et al.*, 2016; Nguyen Chi *et al.*, 2016; Zhong *et al.*, 2017). More than 75 years ago, Lord Adrian working with anesthetized hedgehogs had already demonstrated that the brain produces electrical activity phase-locked to breathing cycles (Adrian, 1942). However, such respiration-driven network oscillations were believed to be restricted to areas involved in olfaction such as the olfactory bulb and piriform cortex (Fontanini *et al.*, 2003; Fontanini & Bower, 2006; Kay *et al.*, 2009). Recent studies have now revealed that RR occurs in many more brain areas than previously thought. Namely, RR has been detected in the hippocampus (Yanovsky *et al.*, 2014; Lockmann *et al.*, 2016; Nguyen Chi *et al.*, 2016), parietal cortex (Zhong *et al.*, 2017), sensory barrel cortex (Ito *et al.*, 2014), and prefrontal cortex (Lockmann *et al.*, 2016; Biskamp *et al.*, 2017; Zhong *et al.*, 2017). Importantly, since RR follows breathing rate, its peak frequency is quite variable and depends on animal species and behavioral state. In rodents, it often assumes values in the delta and theta frequency range (Nguyen Chi *et al.*, 2016; Zhong *et al.*, 2017), which may have precluded its identification as an independent rhythm.

These recent results indicate that respiration-coupled network activity is likely to be a global brain rhythm, which has not been previously recognized as such due to its variable frequency and the usual lack of simultaneous recordings of respiration in LFP studies. In the present work, we sought to expand our previous findings by simultaneously recording respiration and LFPs from 15 regions of freely moving mice. To further differentiate RR from theta, we focused our analysis on two behavioral states in which theta oscillations are prominent: exploration and REM sleep. Our results show that respiration-coupled oscillations occur in parallel with theta in widespread brain regions, including several neocortical areas and subcortical structures such as the thalamus, amygdala, and ventral hippocampus.

## Materials and Methods

Local field potentials (LFPs) were recorded from 15 brain regions of freely moving mice. A total of 54 animals were analyzed during either exploratory behavior or REM sleep. In all animals, respiratory activity was recorded simultaneously with LFPs by either using thermocoupled sensors chronically implanted into the nasal cavity (exploration) or whole-body plethysmography (REM sleep). Further details are provided below.

### Ethics statement

The present study was carried out in agreement with guidelines of the European Science Foundation (2000), the U.S. National Institutes of Health Guide for the Care and Use of Laboratory Animals (2011), and has been approved by the Governmental Supervisory Panel on Animal Experiments of Baden Württemberg at Karlsruhe (35-9185.81/G-84/13 and 35-9185.81/G-115/14).

### Animal Care and Housing Conditions

C57BL/6N mice were purchased at 14 weeks of age from Charles River (Sulzfeld, Germany). Animals were housed in groups of four inside a ventilated Scantainer (Scanbur BK A/S Denmark) on an inverted 12/12-h light/dark cycle (light on at 8:00 p.m.) for a minimum of two weeks. Animals had free access to water and food. Following chronic electrode implantation, mice were housed individually. After finishing recordings, animals were killed with an overdose of isoflurane during brain perfusion.

### Animal Preparation

Fifty four C57BL/6N mice (34 female and 20 male) were used in the present study. Animals weighed between 21 and 40 g and were from 14 to 40 weeks old. For electrode implantation, animals were anesthetized with isoflurane in medical oxygen (4% isoflurane for induction, 1.5–2.5% for maintenance, flow rate: 1 l per min). For analgesia, 0.1 mg/kg of buprenorphine was injected subcutaneously prior to and 8 h after surgery. Anesthetized animals were mounted on a stereotaxic apparatus (Kopf Instruments, Tujunga, CA) with a custom-made inhalation tube. Body temperature was maintained at 38°C by a heating pad (ATC-2000, World Precision Instruments). For monitoring the temperature of nasal air flow, two precision fine bare wire temperature sensors (80 µm diameter, Omega Engineering Inc., Stamford, CT; Part No.: 5TC-TT-KI-40-1M) were implanted into the right and left nasal cavity (11 mm anterior, 0.5 mm lateral). After exposure of the skull, holes of 0.5–1.0 mm in diameter were drilled above the following brain structures: dorsal hippocampus (dHip), ventral hippocampus (vHip), olfactory bulb (OB), prelimbic cortex (PLC), parietal cortex (PAC), anterior cingulate cortex (ACC), somatosensory cortex (SSC), insular cortex (INS), vibrissal area of the motor cortex (vMC), visual cortex (VC), lateral entorhinal cortex (LEC), basal nucleus of amygdala (AMYG), mediodorsal thalamic nucleus (MD), ventroposterior thalamic nucleus (VPL). For stereotaxic coordinates of electrode positions, see Table 1 (Paxinos & Franklin, 2004). Two stainless steel watch screws (1 x 3 mm) over the cerebellum served as ground and reference electrodes. Recording electrode were made of pairs of varnish-insulated tungsten wires (50 µm, glued together) which were implanted into the depths listed in Table 1, with the exception of three surface locations (OBs, PAC and VC), where epidural recordings were performed using watch screws.

**Table 1:** Stereotaxic coordinates and sample size.

| Brain region (Abbreviation) | Coordinates from bregma (mm) |  |  | # mice |
| --- | --- | --- | --- | --- |
|  | AP | ML | DV |  |
| Olfactory bulb, gran. layer (OBd) | 4.50 | 0.80 | 1.30 | 8 |
| Olfactory bulb, surface (OBs) | 4.50 | 0.80 | dura | 8 |
| Anterior cingulate cortex (ACC) | 1.98 | 0.35 | 1.70 | 8 |
| Prelimbic cortex (PLC) | 1.54 | 0.30 | 2.50 | 8 |
| Vibrissal Motor cortex (vMC) | 0.98 | 1.50 | 0.80 | 8 |
| Somatosensory cortex (SSC) | -1.06 | 1.40 | 0.70 | 8 |
| Insular cortex (INS) | -1.06 | 3.70 | 3.50 | 8 |
| Amygdala, central nucl. (AMYG) | -1.06 | 2.20 | 4.70 | 7 |
| Medio-dorsal thalamus (MD) | -1.46 | 0.50 | 3.00 | 8 |
| Ventral posterior thalamus (VPL) | -1.70 | 1.80 | 3.70 | 8 |
| Dorsal hippocampus, CA1 (dHip) | -2.06 | 1.50 | 1.50 | 8 |
| Parietal cortex (PAC) | -2.06 | 1.50 | dura | 8 |
| Visual cortex (VC) | -2.92 | 2.00 | dura | 7 |
| Ventral hippocampus, CA1 (vHip) | -3.16 | 3.00 | 3.70 | 8 |
| Lateral entorhinal Cortex (LEC) | -4.00 | 3.50 | 4.50 | 8 |

AP: anterior-posterior, ML: medio-lateral, DV: dorso-ventral, dura: epidural.

### Electrophysiology

Intracranial monopolar recordings began 6 to 7 days after surgery. Animal’s spontaneous behavior in the home cage was assessed by the video tracking system Ethovision XT 9 (Noldus Information Technology, Wageningen, Netherlands). Movements in the home cage or in the whole-body plethysmograph (EMKA Technologies, S.A.S., France, for details see Jessberger *et al.*, 2016) were detected by 3-D accelerometry. Successive recording sessions of up to 4 h were performed in the animal’s home cage and in the plethysmograph to collect sufficient sections with non-overlapping theta and respiration frequencies (see below). Extracellular signals were filtered (1–500 Hz), amplified (RHA2116 Intan Technologies, LLC), digitized (2.5 kHz) and stored for offline analyses.

### Data analysis

Data were analyzed in MATLAB (The Mathworks Inc., Natick, MA) using built-in and custom-written routines. We focused on periods of exploration and REM sleep recorded in the home cage and plethysmograph, respectively (for detailed descriptions of behavioral staging see Brankačk *et al.*, 2010; Jessberger *et al.*, 2016; Zhong *et al.*, 2017). In both states, theta oscillations and respiration may overlap in frequency. We only used sections where respiration frequency (based on the power spectrum of the respiration signal) and theta frequency (inferred by the LFP power spectrum) were not overlapping. For each animal and region, the analyzed LFP length was fixed at 30 s, obtained by concatenating epochs within periods of exploration and REM sleep with the largest frequency difference between theta and respiration.

#### Spectral and coherence analysis

Power spectral density was calculated by means of the Welch periodogram method using 4-s Hamming windows with 50% overlapping (*pwelch.m* function from the Signal Processing Toolbox). To compute LFP phase coherence to either the respiration or the theta-filtered signal, we used 1-s windows with 50% overlap (*mscohere.m* function from the Signal Processing Toolbox). Filtering into the theta band (5-10 Hz) was obtained by using the *eegfilt.m* function from the EEGLAB toolbox (Delorme & Makeig, 2004); to remove artifacts, the respiration signal was band-pass filtered using cutoff frequencies around its peak in the power spectrum.

#### Power ratio

For a fixed region and behavioral state, the relative power ratio was obtained by subtracting the peak power of RR from the peak power of theta, normalized by their sum:

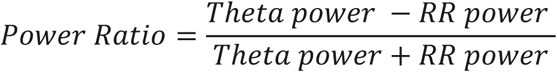

The power ratio varies between -1 and 1; positive values indicate that theta power is stronger than RR power; conversely, negative values indicate that RR power is stronger.

### Histology

After conclusion of the experiments, animals were deeply anesthetized with isoflurane and perfused transcardially with phosphate buffered saline and subsequently with 4% paraformaldehyde (PFA). Brains were carefully dissected, stored in PFA overnight and coronal sections were cut (50 µm), mounted, and stained with cresyl violet. Electrode position was then verified by light microscopy.

### Statistics

For displaying group results (Figures 2 and 4), power and coherence spectra are expressed as means ± SEM over animals. In Figure 5, power ratio values were compared against zero using paired t-tests. Coherence values between LFP and either respiration or the theta reference signal were compared against chance using surrogate-based statistical testing. To that end, surrogate values were obtained by computing coherence spectra between LFPs and reference signals (theta or respiration) from different animals. For each region, the actual distribution of peak values in LFP-respiration or LFP-theta coherence spectra (sample size: # of animals) was compared (unpaired t-test) with the distribution of surrogate coherence values at the corresponding frequency (sample size: [# of animals] X [# of animals -1]).

## Results

We performed multi-site LFP recordings while assessing respiration in a total of 54 freely moving mice (see Table 1 for the exact number of animals per region). Figure 1A shows a representative example of simultaneous recordings of respiration and LFPs from four regions during REM sleep. Respiration was assessed through whole-body plethysmography since nasal thermocouple signals are not fully reliable during sleep (Jessberger *et al.*, 2016). Visual inspection of the traces readily reveals that the parietal cortex LFP exhibited a prominent theta rhythm at ~7 Hz, characteristic of REM sleep. On the other hand, the olfactory bulb and the anterior cingulate cortex displayed a slower LFP rhythm that closely followed nasal respiration at ~3 Hz, which we refer to as the respiration-entrained rhythm (RR). Interestingly, the LFP signal recorded from the insula exhibited both RR and theta activity (compare with the parietal cortex and olfactory bulb LFPs). Figure 1B displays respiratory frequency, LFP power spectra from the respective regions and LFP phase coherence to respiration (red traces) or to a reference theta signal (parietal cortex LFP band-pass filtered at 5-10 Hz; green traces). Notice that all LFPs were coherent with the theta-filtered signal at the theta frequency. At the same time, however, LFPs were also coherent with the respiration signal at the respiration frequency. Therefore, in this example case, all regions exhibited both theta and a slower respiration-coupled rhythm during REM sleep, albeit with different relative magnitudes.

**Figure 1.**
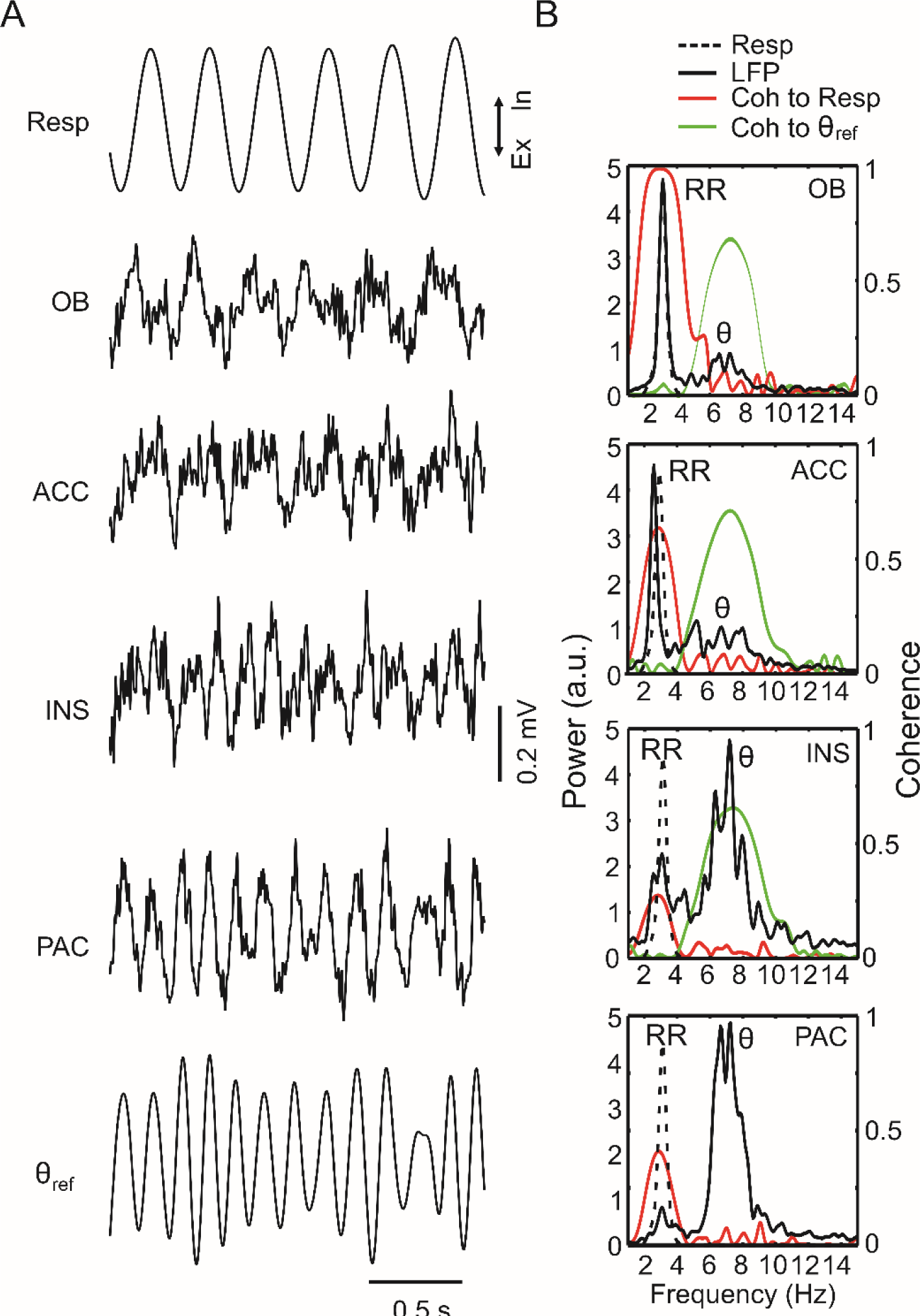
Parallel occurrence of theta and respiration-entrained rhythms in 4 regions of the mouse brain during REM sleep. (A) Traces show two seconds of LFP signals from the olfactory bulb (OB), anterior cingulate cortex (ACC), insular cortex (INS) and parietal cortex (PAC) along with respiration (Resp) and the theta-filtered component of the PAC LFP (θ_ref_). Ex: expiration; In: inspiration. (B) Power spectra of LFPs (solid black lines) and Resp (dotted black lines; same in all panels) plotted along with coherence (Coh) spectra between LFP and Resp (red lines) and between LFP and θ_ref_ (green lines). Notice, in addition to theta activity (θ), power and coherence peaks at the same frequency as Resp, which indicates the presence of a respiration-entrained LFP rhythm (RR). Results obtained from simultaneous recordings in a representative animal using 30-s of concatenated REM sleep epochs.

Figure 2 shows that similar results hold at the group level during REM sleep, and extends to all fifteen brain regions analyzed (mean RR frequency: 2.97 ± 0.06 Hz; mean theta frequency: 7.10 ± 0.07 Hz). In general, RR was most prominent in frontal regions such as the olfactory bulb, prelimbic cortex, and anterior cingulate cortex. Nevertheless, RR could also be detected at a smaller magnitude in diverse other areas such as the thalamus, ventral hippocampus, and lateral entorhinal cortex. Similarly, theta oscillations could also be detected in several regions, though with lower amplitude in the frontal regions where RR activity prevailed. Of note, phase coherence between LFP and either respiration or the theta reference signal was significantly higher than chance in all recorded regions, irrespectively of the magnitude of RR and theta (Figure S1 and Tables S1 and S2). Therefore, we conclude that RR and theta may simultaneously occur in widespread regions of the mouse brain during REM sleep. While they are not mutually exclusive, RR is most noticeable at frontal regions while theta dominates more posteriorly.

**Figure 2.**
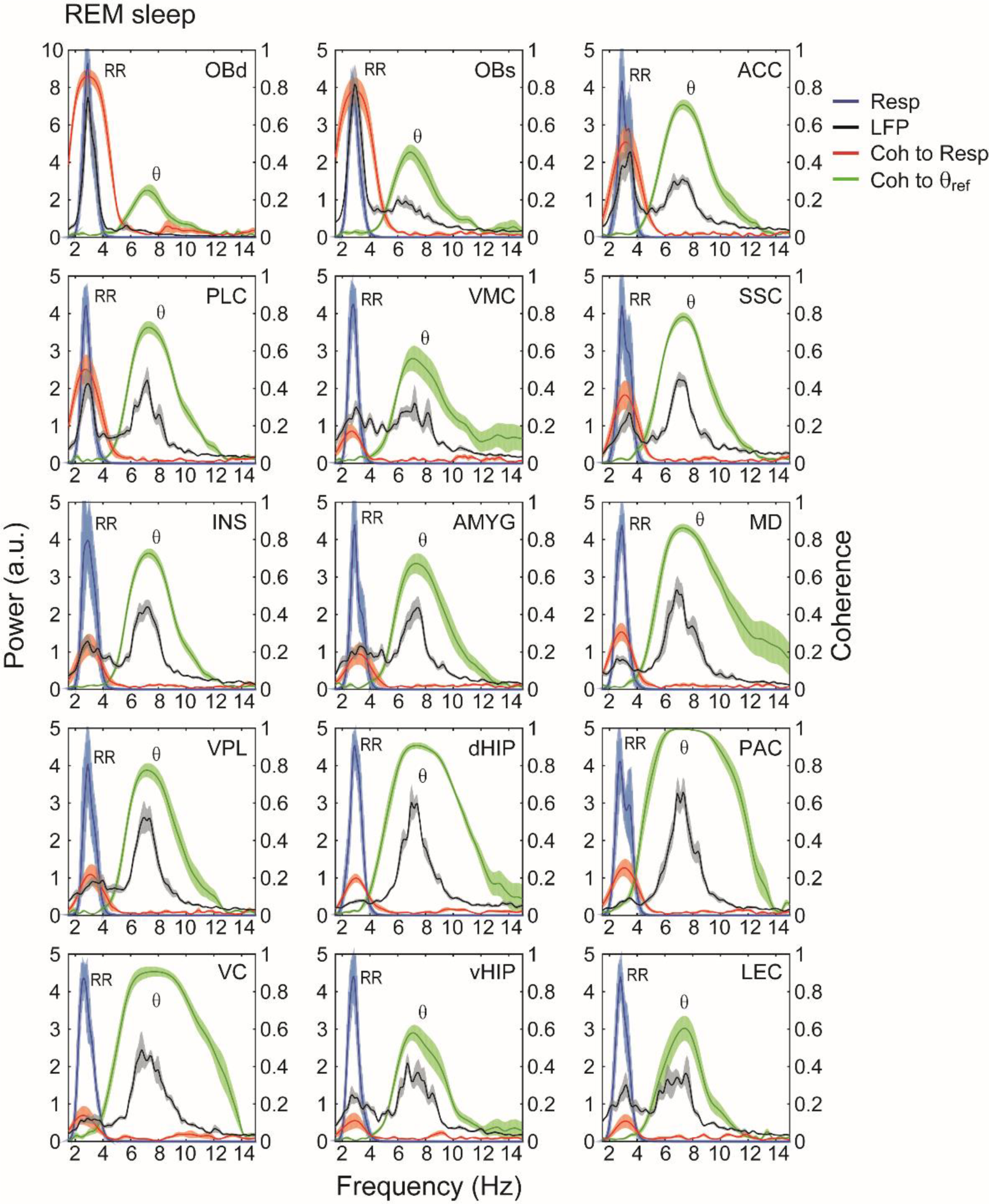
Parallel occurrence of theta (θ) and respiration-entrained LFP rhythm (RR) throughout the mouse brain during REM sleep. Panels show mean power spectral densities of local field potentials (LFP) (black lines) and their coherence (Coh) to respiration (Resp) (red lines) or theta (θ_ref_) (green lines) in fifteen brain regions during REM sleep (30 seconds of concatenated data per animal). Shades represent ± SEM. The reference theta-filtered signal was taken from either the dorsal hippocampus or the parietal cortex. Respiration was assessed through whole-body plethysmography. Power spectra of respiration are also shown (blue lines). Notice LFP power peaks as well as LFP-Resp coherence peaks at the respiration frequency in all regions. OBd: deep olfactory bulb (granular cell layer); OBs: surface of olfactory bulb; ACC: anterior cingulate cortex; PLC: prelimbic cortex; VMC: vibrissal area of motor cortex; SSC: somatosensory cortex; INS: insular cortex; AMYG: amygdala; MD: mediodorsal thalamus; VPL: ventral posterior lateral thalamus; dHip: dorsal hippocampus; PAC: parietal cortex; VC: visual cortex; vHip: ventral hippocampus; LEC: lateral entorhinal cortex.

We next analyzed awake periods in which the animal actively explored the home cage, a behavior that induces robust theta oscillations (Zhong *et al.*, 2017). Nasal respiration was tracked using temperature sensors implanted into the nostrils. It should be noted that during locomotion and exploration the breathing rate of mice may be the same or even faster than theta frequency (Wesson *et al.*, 2008; Nguyen Chi *et al.*, 2016; Zhong *et al.*, 2017). As previously argued (Yanovsky *et al.*, 2014; Nguyen Chi *et al.*, 2016; Zhong *et al.*, 2017), it can, therefore, be difficult to disentangle both rhythms. Here we opted to focus on exploration periods in which animals breathed faster than theta, inferred by two clearly separate signals in power spectra. Figure 3A shows example traces of LFPs and respiration in a representative animal during such a period. Notice prominent theta oscillations at ~8.5 Hz in the dorsal hippocampus. On the other hand, the olfactory bulb LFP exhibited faster oscillations at ~12 Hz that were clearly locked to nasal respiration, thus characterizing RR activity. Interestingly, as shown in Figure 3B, the power spectrum of the LFP recorded from the prelimbic cortex revealed two peaks, one at the same frequency as the hippocampal power peak and corresponding to theta oscillations and the other at the same frequency as respiration and corresponding to RR. Notice back in Figure 3A that it is very difficult to infer the existence of either rhythm solely by visual inspection of the prelimbic cortex LFP trace. This is because the simultaneous presence of both theta and RR leads to alternating effects of constructive and destructive interferences that give rise to frequency beating (see Figure 3 in Yanovsky *et al.*, 2014). Finally, although not apparent upon visual inspection (Figure 3A), the power and coherence spectra reveal that RR was also present in the mediodorsal nucleus of the thalamus and in the dorsal hippocampus, but at a much lower magnitude (Figure 3B). Therefore, and similarly to the example in Figure 1, all recorded regions simultaneously exhibited RR and theta during exploration.

**Figure 3.**
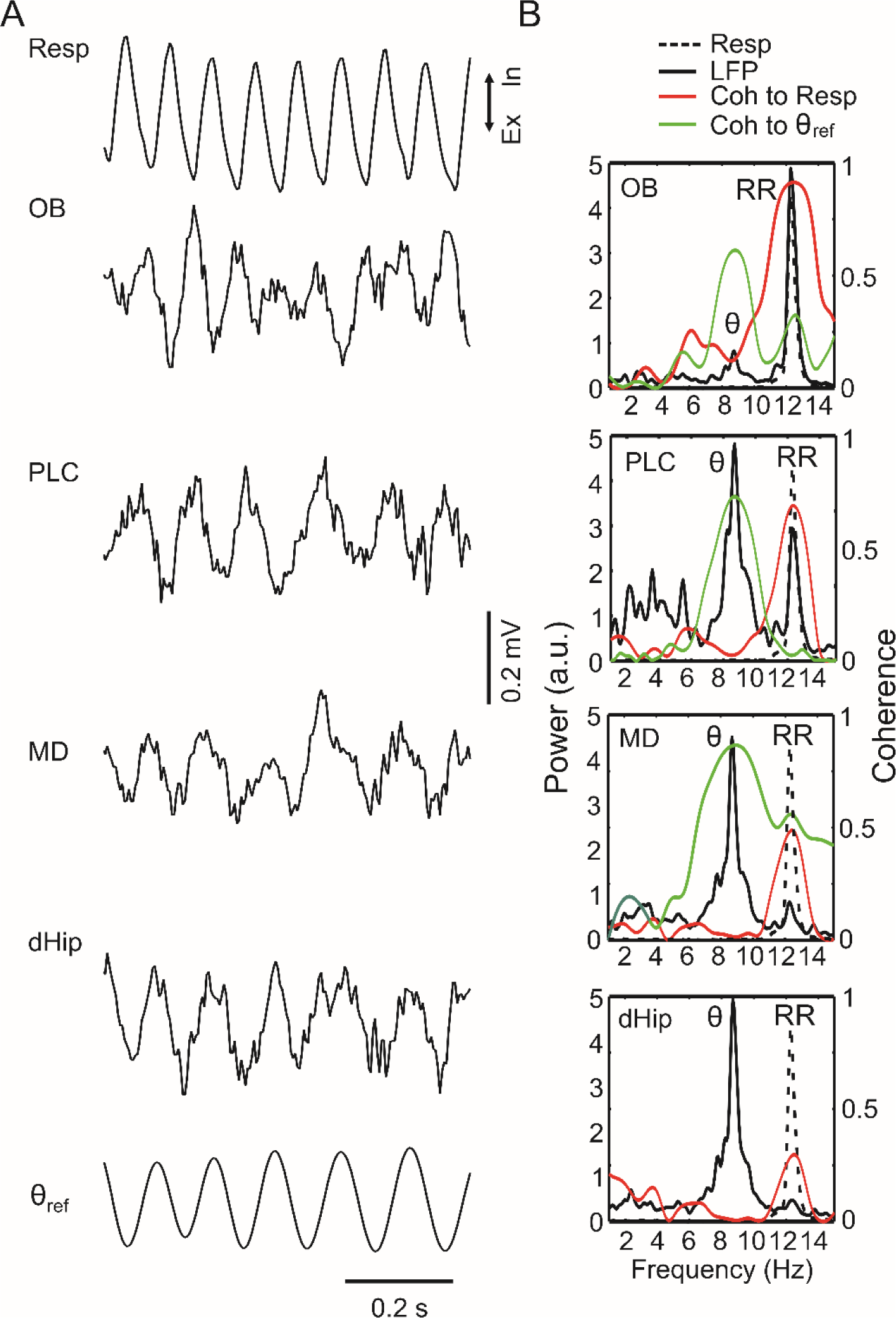
Theta and respiration-entrained rhythms in a representative animal during exploration. (A,B) Panels as in Figure 1A,B. The reference theta signal (θ_ref_) was obtained from the dorsal hippocampus. In B, we analyzed 30-s of concatenated epochs in which the animal explored the environment while breathing at a rate faster than theta. OB: olfactory bulb; PLC: prelimbic cortex; MD: medio-dorsal thalamus; dHip: dorsal hippocampus; RR: respiration-entrained rhythm; Ex: expiration; In: inspiration.

Figure 4 shows group results of LFP power spectra and LFP coherence to theta and respiration for 15 brain regions recorded while animals were engaged in exploration with respiration faster than theta (mean RR frequency: 11.03 ± 0.12 Hz; mean theta frequency: 8.50 ± 0.09 Hz). Notice parallel occurrence of both rhythms in several regions. Consistently, peak coherence values between LFPs and either reference signal (respiration or theta) were significantly higher than chance in all recorded regions during exploration (Figure S2 and Tables S3 and S4). Finally, Figure 5 provides group data for the spatial distribution of LFP coherence to respiration or theta, as well as of the relative power between theta and RR within each of the 15 brain regions. In the bar graphs, a relative power ratio of 1 denotes exclusive theta activity while -1 denotes exclusive RR activity; a relative power ratio of 0 means that both rhythms had the same magnitude. Notice similar distributions of RR and theta activities during REM sleep and exploration: in either behavioral state, RR was most prominent in frontal regions while theta prevailed in more posterior regions, with no region exclusively exhibiting only theta or RR. In all, our results show that not only theta but also RR constitutes a global brain rhythm.

**Figure 4.**
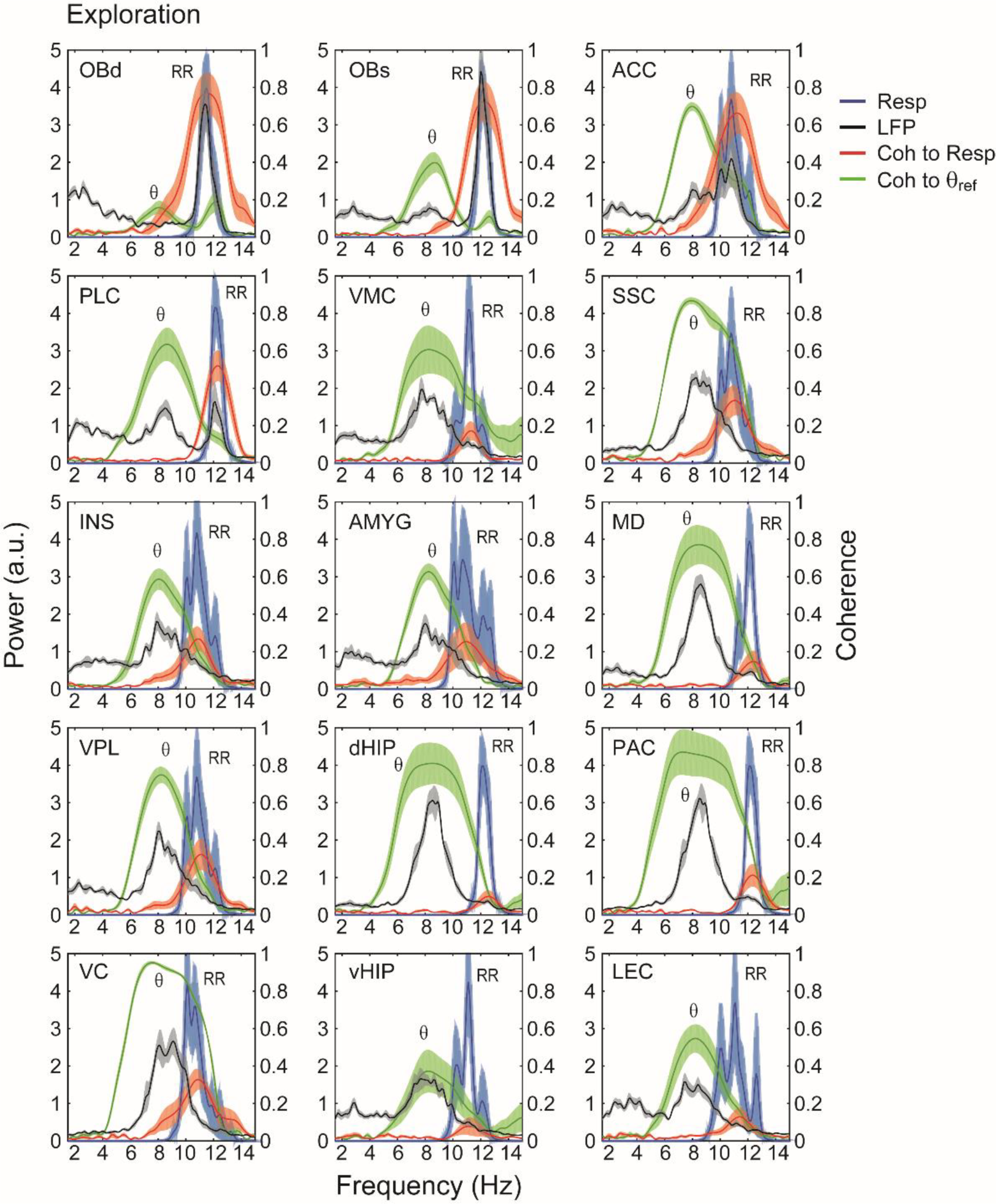
Parallel occurrence of theta (θ) and respiration-entrained LFP rhythm (RR) throughout the mouse brain during exploration. Panels show the same as in Figure 2. Each sample consisted of 30 seconds of concatenated data obtained during exploration with respiration faster than theta. The reference theta-filtered signal was taken from either the dorsal hippocampus or the parietal cortex. Respiration was assessed through thermocouples in the nasal cavity. OBd: deep olfactory bulb (granular cell layer); OBs: surface of olfactory bulb; ACC: anterior cingulate cortex; PLC: prelimbic cortex; VMC: vibrissal area of motor cortex; SSC: somatosensory cortex; INS: insular cortex; AMYG: amygdala; MD: mediodorsal thalamus; VPL: ventral posterior lateral thalamus; dHip: dorsal hippocampus; PAC: parietal cortex; VC: visual cortex; vHip: ventral hippocampus; LEC: lateral entorhinal cortex.

**Figure 5.**
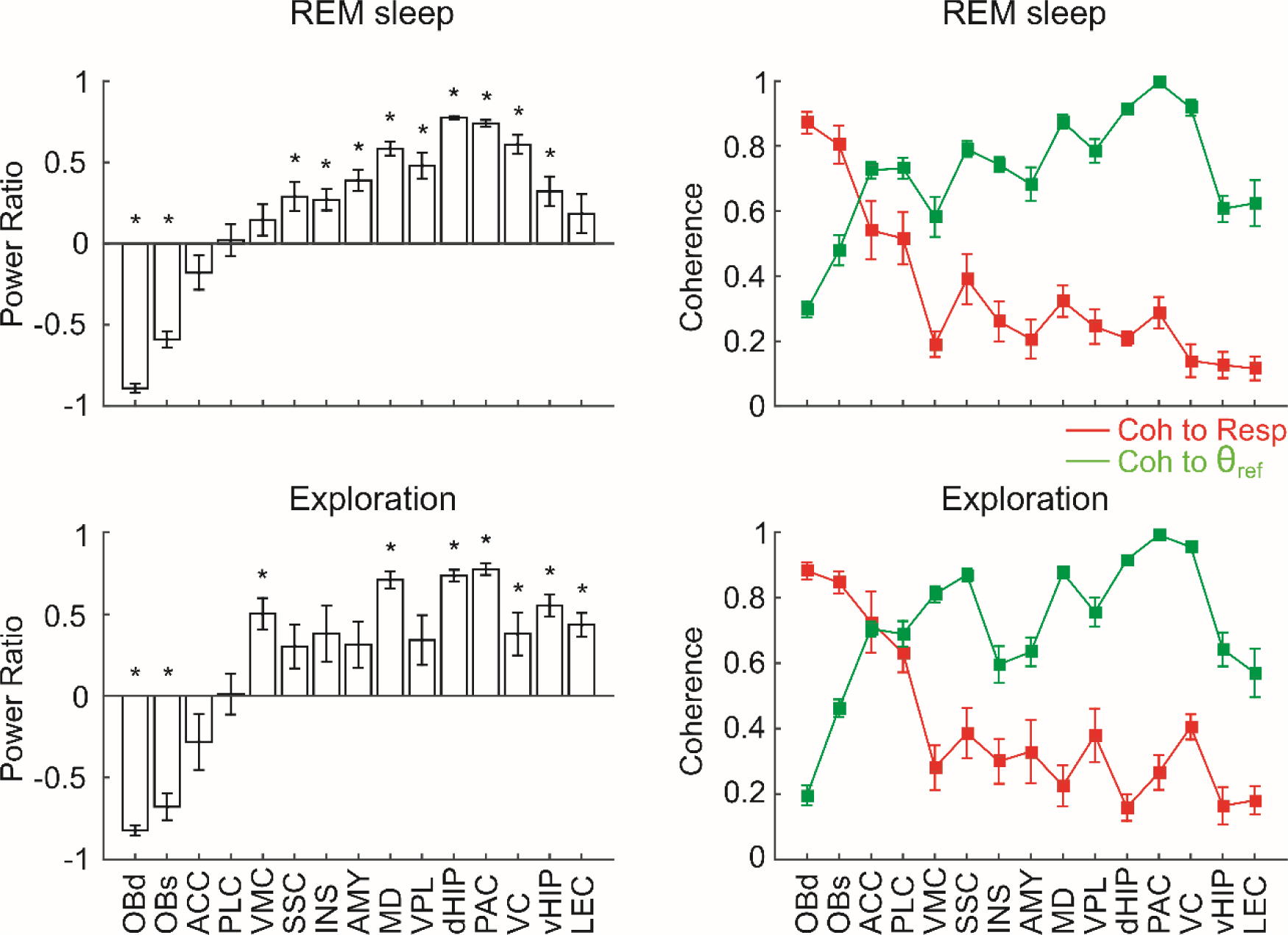
Spatial distribution of the respiration-entrained rhythm (RR) and theta (θ) activity during REM sleep and exploration. The bar graphs on the left show the mean relative power ratio (±SEM) between θ and RR for each recorded region. Note that no brain region exhibits θ alone (power ratio = 1.0) or RR alone (power ratio = -1). *p<0.05 compared to 0 (paired t-tests). The right panels show mean LFP coherence to respiration (red) or to theta (green) (±SEM). The reference theta-filtered signal was taken from either the dorsal hippocampus or the parietal cortex. Notice similar distributions of theta and RR activity during REM sleep and exploration. OBd: deep olfactory bulb (granular cell layer); OBs: surface of olfactory bulb; ACC: anterior cingulate cortex; PLC: prelimbic cortex; VMC: vibrissal area of motor cortex; SSC: somatosensory cortex; INS: insular cortex; AMY: amygdala; MD: mediodorsal thalamus; VPL: ventral posterior lateral thalamus; dHip: dorsal hippocampus; PAC: parietal cortex; VC: visual cortex; vHip: ventral hippocampus; LEC: lateral entorhinal cortex.

## Discussion

We have simultaneously tracked nasal respiration along with multisite LFP recordings in freely moving mice during two behavioral states classically associated with theta oscillations. For each region, we computed phase coherence spectra between its LFP and either respiration or a reference theta signal. As in previous work, we inferred the existence of a respiration-locked network rhythm if (1) the LFP exhibited a power peak at the same frequency as respiration, and (2) if the LFP coherence with respiration peaked at this frequency. We found that the amplitude of RR was highest in frontal regions, such as the olfactory bulb, prelimbic cortex, and anterior cingulate cortex. Nevertheless, a power peak at the same frequency as respiration and coherent with it was also apparent at more ventral and posterior regions, such as the visual cortex, amygdala, ventral hippocampus and lateral entorhinal cortex. In all, our results provide evidence that respiration-locked network oscillations can be detected in several brain regions, including cortical and subcortical structures, where they may occur concomitantly with theta oscillations.

Mice and rats can breathe as slow as 1 Hz during quiet states and as fast as 14-15 Hz during running, exploration and sniffing (Wesson *et al.*, 2008; Rojas-Líbano *et al.*, 2014). Therefore, the respiration-coupled LFP rhythm escapes any simple frequency-based definition. Its peak frequency can be the same as that of distinct network oscillations: slow thalamo-cortical oscillations as observed during sleep (~0.3-1 Hz; Crunelli & Hughes, 2010), delta oscillations (1-5 Hz; Lakatos *et al.*, 2005), theta oscillations (5-10 Hz; Nguyen Chi *et al.*, 2016), neocortical mu and alpha rhythms (8-12 Hz; Tort *et al.*, 2010), or even oscillations in the low beta range (12-20 Hz; Haenschel *et al.*, 2000). In particular, in this work we showed examples of RR at 2-4 Hz during REM sleep, i.e. slower than theta (Figures 1 and 2), and at 10-14 Hz during exploration, i.e. faster than theta (Figures 3 and 4). We intentionally avoided selecting epochs in which breathing rate occurred at theta frequency to more clearly demonstrate the distinction between the two rhythms, but these periods often occur during active behaviors. Moreover, RR tends to have larger amplitude when animals breathe at lower frequencies (Jessberger *et al.*, 2016).

It should be noted that previous studies have been referring to RR in the olfactory bulb and piriform cortex as “olfactory theta” (Kay & Freeman, 1998; Kay *et al.*, 2009; Rojas-Líbano *et al.*, 2014). We do not favor such a nomenclature because (1) it may give the idea of a mechanistic link between RR and hippocampal theta oscillations, while we have shown that the two rhythms are independent (Yanovsky *et al.*, 2014; Lockmann *et al.*, 2016; Nguyen Chi *et al.*, 2016); and (2) it may mask the fact that olfactory areas may exhibit two peaks within the theta band, one due to the classical theta rhythm and another due to RR (Nguyen Chi *et al.*, 2016; Zhong *et al.*, 2017). Moreover, (3) it may also mask the fact that RR can have peak frequencies much below or much higher than the traditional theta frequency range. Again, given its variability in peak frequency, we consider that a simple frequency-based definition would not be proper for this rhythm.

Our results indicate that respiration-coupled oscillations are a global brain rhythm. This raises the possibility that previous research on LFP oscillations may have been “contaminated” by RR, which was not recognized due to the lack of simultaneous recordings of respiration in the experiments. As an example, we note that whether the slow oscillations observed during deep sleep and anesthesia (“up-and-down” transitions; Clement *et al.*, 2008) would couple or not to respiration has been disputed, with evidence for (Fontanini *et al.*, 2003) and against (Viczko *et al.*, 2014). We have recently solved this debate by showing that there are two oscillations of nearby frequency (0.3-1.5 Hz) during these states, one corresponding to the up-and-down transitions and the other to RR (Lockmann *et al.*, 2016). Given the proximity in peak frequency, some studies likely confounded the latter with the former (see Fontanini & Bower, 2006). Similarly, Ito et al. (2014) have recently described that “delta” oscillations in the barrel cortex couple to respiration. While our results corroborate the existence of such respiration-coupled oscillations in sensory cortices, we would be less inclined to conclude that all delta-frequency activity in the barrel cortex is due to respiration, as apparently genuine delta oscillations occur during sleep in several neocortical regions of rodents (Watson *et al.*, 2016). As another example, we also believe that the “slow theta” oscillations in the striatum that have been shown to vary in peak frequency during T-maze traversals and to modulate 80-120 Hz oscillations (Tort *et al.*, 2008) do most likely correspond to RR (see Zhong *et al.*, 2017). The required re-classification of some slow network oscillations does not, of course, argue against the importance of previous reports. Indeed, it is likely that respiration-coupled oscillations provide a mechanism for binding different networks and neuronal ensembles into a common regime, and thus fulfill similar behavioral and cognitive functions as other rhythms in the same frequency domain. This may even extend to humans where recent work shows the presence, and cognitive relevance, of LFP activity entrained by respiration in hippocampal and amygdala circuits (Zelano *et al.*, 2016).

Importantly, the current and former results show that respiration-coupled oscillations are particularly prominent in the medial prefrontal cortex (mPFC) (Lockmann *et al.*, 2016; Biskamp *et al.*, 2017; Zhong *et al.*, 2017). Since RR also modulates gamma (Biskamp *et al.*, 2017; Zhong *et al.*, 2017), we suspect that the coupling between “theta” and gamma recently reported in the mPFC (Zhang *et al.*, 2016; Pavlovsky *et al.*, 2017) actually corresponds to respiration-gamma coupling. Furthermore, theta-frequency oscillations were previously shown to synchronize activity in the mPFC and in the ventral hippocampus during anxiety (Adhikari *et al.*, 2010). Since our results show that both theta and RR exist in these regions, such experiments warrant being revisited with simultaneous recordings of respiration. Finally, we also suggest that the “4-Hz oscillation” described to link mPFC, ventral tegmental area and hippocampus during working memory (Fujisawa & Buzsáki, 2011; Guise & Shapiro, 2017; see also Roy *et al.*, 2017), and to coordinate mPFC and amygdala networks during fear learning (Dejean *et al.*, 2016; Karalis *et al.*, 2016), may correspond to respiration-coupled oscillations.

While our work shows that RR can be detected in LFPs from widespread regions of the mouse brain, from prefrontal to visual cortices, it remains to be determined whether RR is generated de novo in each of these regions or, alternatively, if it is volume conducted. We note that a similar issue applies to theta oscillations: even though theta has been hypothesized to play functional roles in several regions, its local origin in the recorded region has been seldom demonstrated (Jones & Wilson, 2005; DeCoteau *et al.*, 2007; Young & McNaughton, 2009; Adhikari *et al.*, 2010; Benchenane *et al.*, 2010; O’Neill *et al.*, 2013; but see Sirota *et al.*, 2008). Local generation of an oscillation is typically inferred by either bipolar recordings, current-source density analysis (CSD), or modulation of unit activity in the recorded region. In addition to its well-accepted generation in the olfactory bulb, RR has also been demonstrated to be generated in the hippocampus, where CSD analysis revealed maximal activity in the dentate gyrus (Yanovsky *et al.*, 2014; Lockmann *et al.*, 2016). RR was also shown to modulate spiking probability in the hippocampus, somatosensory cortex, parietal cortex, and prefrontal cortex, suggestive of a local origin in these areas (Ito *et al.*, 2014; Yanovsky *et al.*, 2014; Nguyen Chi *et al.*, 2016; Biskamp *et al.*, 2017; Zhong *et al.*, 2017). However, in several other regions the local origin of RR and theta have yet to be experimentally determined.

In summary, our current results indicate that respiration-coupled oscillations are a global brain rhythm, and, as such, it should be put at center stage of research along with theta oscillations. The widespread presence of RR was probably not recognized previously due to the usual lack of simultaneous recordings of respiration along with LFPs. We believe that previous functions attributed to oscillations such as slow oscillations, delta, “4-Hz oscillations” and theta could be due to RR activity.

## Acknowledgments

This work was supported by the Deutsche Forschungsgemeinschaft (SFB 636/B06; SFB 1134/A01; Dr 326/10-1), Bundesministerium für Bildung und Forschung (Bernstein Center for Computational Neurosciences, No. 01GQ1003A; German-Brazil Cooperation grant: No. 01DN12098), the Brazilian National Council for Scientific and Technological Development (CNPq), the Brazilian Coordination for the Improvement of Higher Education Personal (CAPES), and the Alexander von Humboldt Foundation.

## Conflict of interest

The authors declare no competing financial interests.

## Author contributions

A.B.L.T., J.B and A.D. conceived the study. S.P., J.J. and Y.Y. performed experiments. A.B.L.T. and J.B. analyzed the data. All authors discussed results. A.B.L.T., J.B and A.D. wrote the manuscript.

## Data accessibility

Data are available from the corresponding author upon reasonable request.

## Abbreviations

ACC: anterior cingulate cortex
AMYG: amygdala
CSD: current-source density
dHip: dorsal hippocampus
EEG: electroencephalogram
fMRI: functional magnetic resonance imaging
INS: insular cortex
LEC: lateral entorhinal cortex
LFP: local field potential
MD: mediodorsal thalamus
OBd: deep olfactory bulb
OBs: surface of olfactory bulb
PAC: parietal cortex
PFA: paraformaldehyde
PLC: prelimbic cortex
RR: respiration-entrained rhythm
SSC: somatosensory cortex
VC: visual cortex
vHip: ventral hippocampus
vMC: vibrissal area of motor cortex
VPL: ventral posterior lateral thalamus

## Supporting Information

### Abbreviations

ACC: anterior cingulate cortex
AMYG: amygdala
Coh: coherence
dHip: dorsal hippocampus
INS: insular cortex
LEC: lateral entorhinal cortex
LFP: local field potential
MD: mediodorsal thalamus
OBd: deep olfactory bulb
OBs: surface of olfactory bulb
PAC: parietal cortex
PLC: prelimbic cortex
Resp: respiration
RR: respiration-entrained rhythm
SSC: somatosensory cortex
Surrog: surrogate
θ_ref_: theta reference signal
VC: visual cortex
vHip: ventral hippocampus
vMC: vibrissal area of motor cortex
VPL: ventral posterior lateral thalamus

---

**Figure S1.**
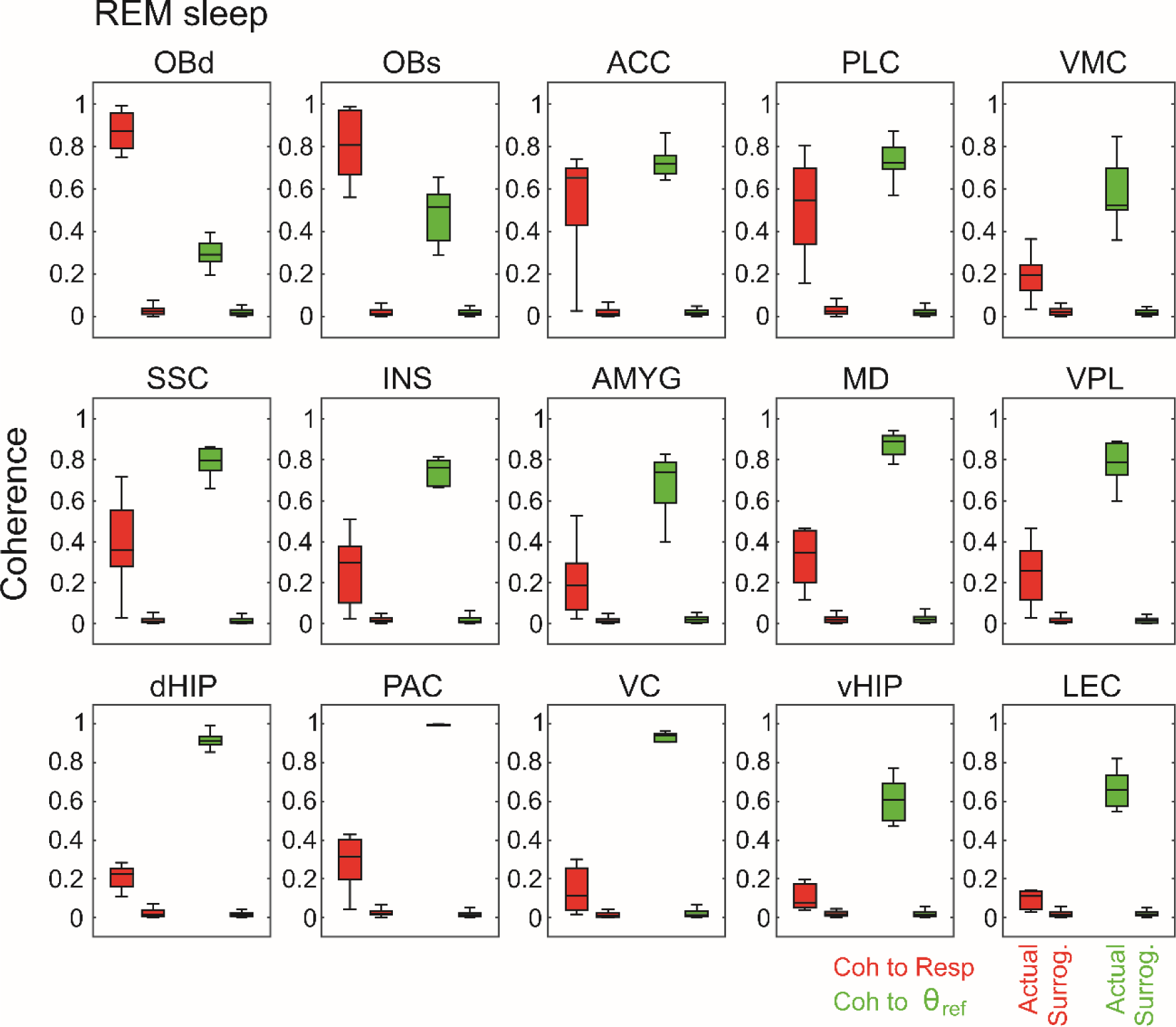
The levels of LFP coherence to either respiration or theta is greater than chance in all recorded regions during REM sleep. Panels show boxplot distributions of the peak coherence values for actual and surrogated data. For each region, surrogated values were obtained by computing phase coherence between the LFP recorded from one animal with the reference signal (respiration or theta) recorded from another animal (see Materials and Methods). This surrogate procedure was performed for all possible pairwise combinations (e.g., for a region with 8 mice recorded, the number of surrogate samples is 8x7=56). The actual and surrogate distributions statistically differ in all recorded regions for both LFP-respiration (red) and LFP-theta coherence (green). See Tables S1 and S2 for statistical analysis.

**Figure S2.**
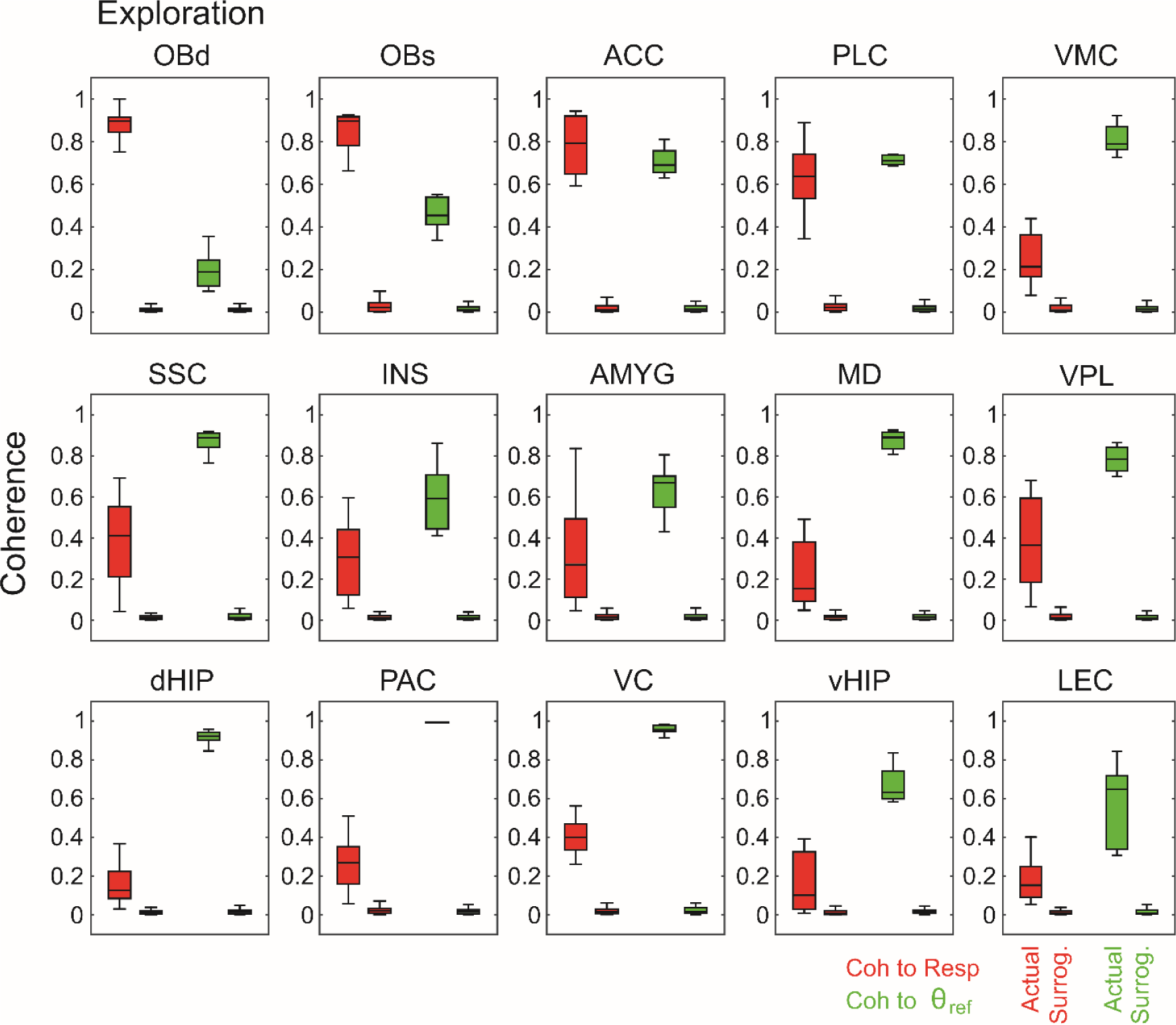
The levels of LFP coherence to either respiration or theta is greater than chance in all recorded regions during exploration. Panels show the same as in Figure S1, but for exploration. See Tables S3 and S4 for statistical analysis.

**Table S1:** LFP-Resp coherence during REM sleep

| Region | Mean Coherence | | 99% Confidence Interval of $\Delta\text{Coh}$ | df | *t* | *P* |
| --- | --- | --- | --- | --- | --- | --- |
|  | Actual | Surrogate |  |  |  |  |
| OBd | 0.871 | 0.034 | 0.793-0.882 | 62 | 50.3 | 0 |
| OBs | 0.803 | 0.019 | 0.726-0.842 | 62 | 35.7 | 4.4E-43 |
| ACC | 0.541 | 0.021 | 0.432-0.607 | 62 | 15.8 | 2.0E-23 |
| PLC | 0.516 | 0.034 | 0.400-0.564 | 62 | 15.6 | 4.4E-23 |
| VMC | 0.191 | 0.026 | 0.115-0.214 | 47 | 8.9 | 1.3E-11 |
| SSC | 0.391 | 0.018 | 0.297-0.448 | 62 | 13.1 | 1.4E-19 |
| INS | 0.261 | 0.021 | 0.179-0.301 | 62 | 10.4 | 3.4E-15 |
| AMYG | 0.207 | 0.020 | 0.128-0.246 | 62 | 8.4 | 8.6E-12 |
| MD | 0.323 | 0.026 | 0.244-0.349 | 62 | 14.9 | 3.3E-22 |
| VPL | 0.244 | 0.016 | 0.175-0.280 | 62 | 11.5 | 4.7E-17 |
| dHIP | 0.209 | 0.022 | 0.159-0.214 | 62 | 18.0 | 2.5E-26 |
| PAC | 0.288 | 0.028 | 0.208-0.311 | 62 | 13.4 | 5.6E-20 |
| VC | 0.140 | 0.020 | 0.056-0.184 | 34 | 5.1 | 1.2E-05 |
| vHIP | 0.126 | 0.020 | 0.065-0.148 | 62 | 6.9 | 3.5E-09 |
| LEC | 0.116 | 0.017 | 0.058-0.139 | 47 | 6.5 | 4.3E-08 |

**Table S2:**
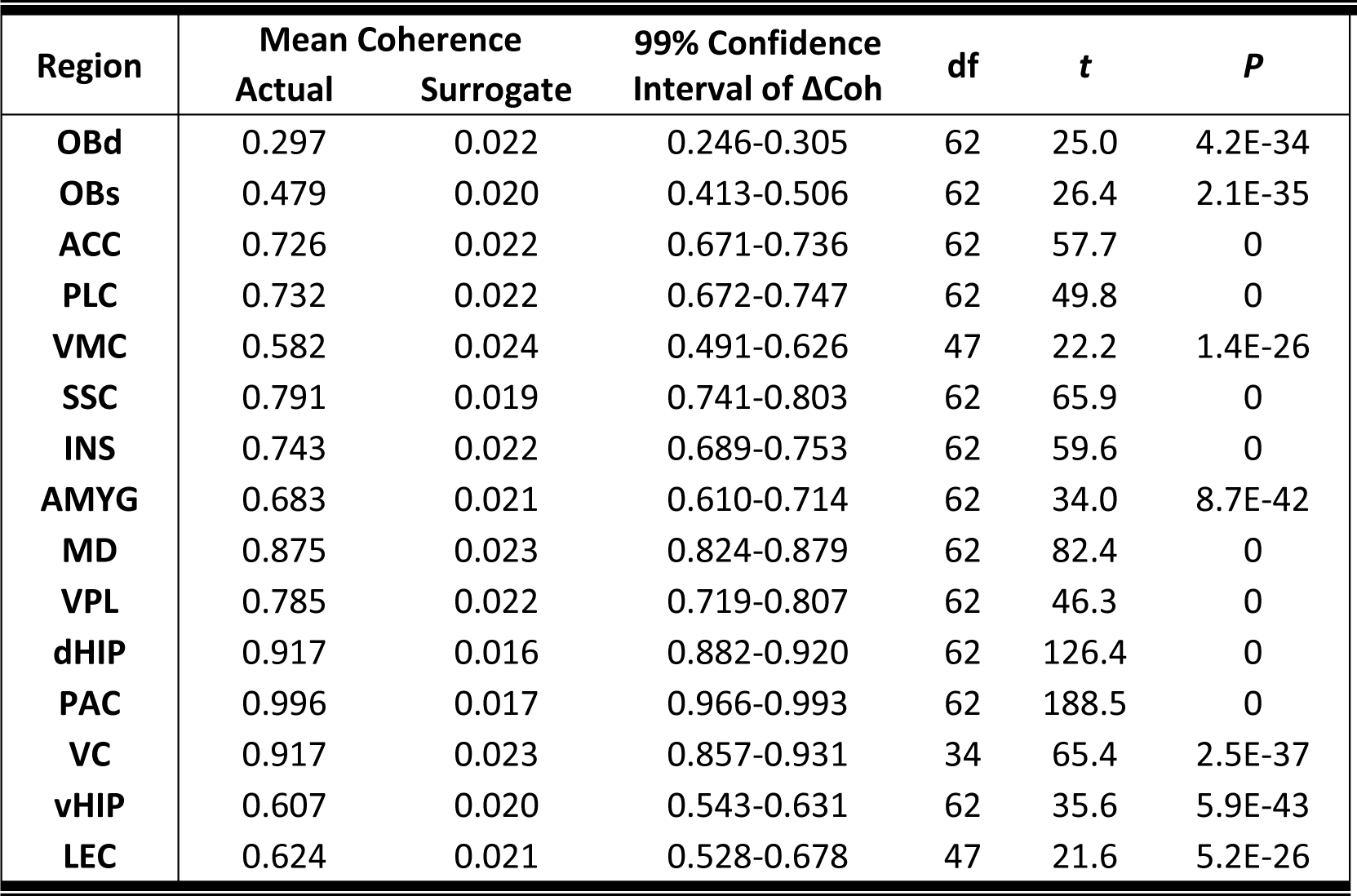
LFP-Theta coherence during REM sleep

**Table S3:**
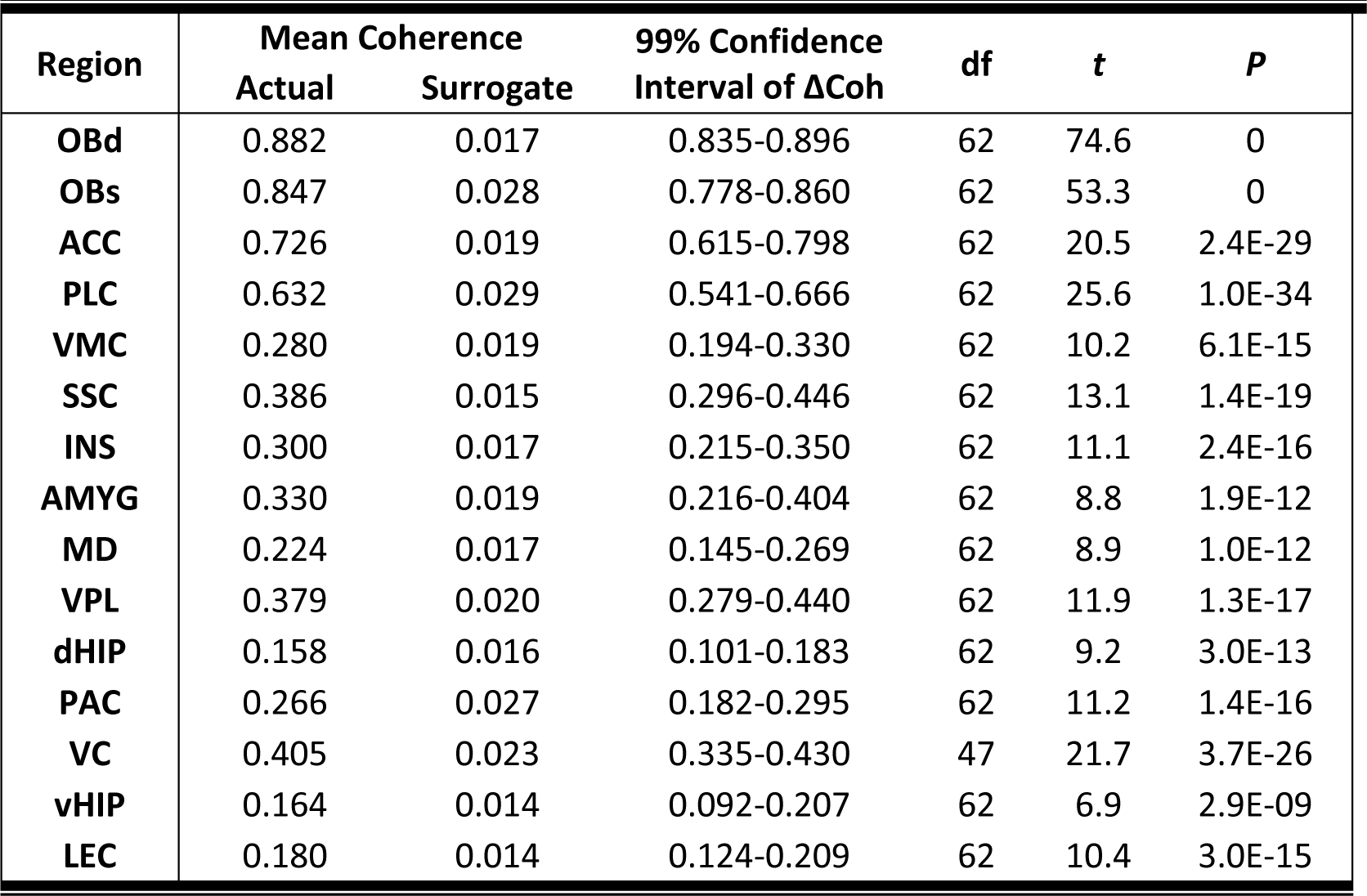
LFP-Resp coherence during Exploration

| Region | Mean Coherence | | 99% Confidence Interval of $\Delta\text{Coh}$ | df | *t* | *P* |
| --- | --- | --- | --- | --- | --- | --- |
|  | Actual | Surrogate |  |  |  |  |
| OBd | 0.882 | 0.017 | 0.835-0.896 | 62 | 74.6 | 0 |
| OBs | 0.847 | 0.028 | 0.778-0.860 | 62 | 53.3 | 0 |
| ACC | 0.726 | 0.019 | 0.615-0.798 | 62 | 20.5 | 2.4E-29 |
| PLC | 0.632 | 0.029 | 0.541-0.666 | 62 | 25.6 | 1.0E-34 |
| VMC | 0.280 | 0.019 | 0.194-0.330 | 62 | 10.2 | 6.1E-15 |
| SSC | 0.386 | 0.015 | 0.296-0.446 | 62 | 13.1 | 1.4E-19 |
| INS | 0.300 | 0.017 | 0.215-0.350 | 62 | 11.1 | 2.4E-16 |
| AMYG | 0.330 | 0.019 | 0.216-0.404 | 62 | 8.8 | 1.9E-12 |
| MD | 0.224 | 0.017 | 0.145-0.269 | 62 | 8.9 | 1.0E-12 |
| VPL | 0.379 | 0.020 | 0.279-0.440 | 62 | 11.9 | 1.3E-17 |
| dHIP | 0.158 | 0.016 | 0.101-0.183 | 62 | 9.2 | 3.0E-13 |
| PAC | 0.266 | 0.027 | 0.182-0.295 | 62 | 11.2 | 1.4E-16 |
| VC | 0.405 | 0.023 | 0.335-0.430 | 47 | 21.7 | 3.7E-26 |
| vHIP | 0.164 | 0.014 | 0.092-0.207 | 62 | 6.9 | 2.9E-09 |
| LEC | 0.180 | 0.014 | 0.124-0.209 | 62 | 10.4 | 3.0E-15 |

**Table S4:** LFP-Theta coherence during Exploration

| Region | Mean Coherence | | 99% Confidence Interval of $\Delta\text{Coh}$ | df | *t* | *P* |
| --- | --- | --- | --- | --- | --- | --- |
|  | Actual | Surrogate |  |  |  |  |
| OBd | 0.196 | 0.015 | 0.149-0.212 | 62 | 15.2 | 1.3E-22 |
| OBs | 0.462 | 0.018 | 0.412-0.476 | 62 | 36.7 | 8.7E-44 |
| ACC | 0.705 | 0.019 | 0.657-0.715 | 62 | 62.9 | 0 |
| PLC | 0.689 | 0.020 | 0.629-0.711 | 62 | 43.2 | 0 |
| VMC | 0.811 | 0.018 | 0.764-0.821 | 62 | 73.9 | 0 |
| SSC | 0.871 | 0.019 | 0.824-0.878 | 62 | 83.9 | 0 |
| INS | 0.596 | 0.017 | 0.522-0.635 | 62 | 27.3 | 2.7E-36 |
| AMYG | 0.635 | 0.020 | 0.568-0.661 | 62 | 35.1 | 1.4E-42 |
| MD | 0.877 | 0.023 | 0.822-0.886 | 62 | 70.6 | 0 |
| VPL | 0.756 | 0.018 | 0.693-0.784 | 62 | 42.8 | 0 |
| dHIP | 0.917 | 0.018 | 0.879-0.917 | 62 | 125.6 | 0 |
| PAC | 0.992 | 0.017 | 0.964-0.987 | 62 | 220.2 | 0 |
| VC | 0.956 | 0.024 | 0.909-0.954 | 47 | 111.0 | 0 |
| vHIP | 0.643 | 0.018 | 0.574-0.676 | 62 | 32.5 | 1.3E-40 |
| LEC | 0.570 | 0.019 | 0.477-0.626 | 62 | 19.7 | 2.1E-28 |

